# Identification of *Fusarium solani* f. sp. *pisi* (*Fsp*) responsive genes in *Pisum sativum*

**DOI:** 10.1101/2020.05.12.091892

**Authors:** Bruce A. Williamson-Benavides, Richard Sharpe, Grant Nelson, Eliane T. Bodah, Lyndon D. Porter, Amit Dhingra

**Author notes:** Correspondence: Amit Dhingra.

## Abstract

*Pisum sativum* (pea) is rapidly emerging as an inexpensive and major contributor to the plant-derived protein market. Due to its nitrogen-fixation capability, short life cycle, and low water usage, pea is a useful cover-and-break crop that requires minimal external inputs. It is critical for sustainable agriculture and indispensable for future food security. Root rot in pea, caused by the fungal pathogen *Fusarium solani* f. sp. *pisi* (*Fsp*), can result in a 15-60% reduction in yield. It is urgent to understand the molecular basis of *Fsp* interaction in pea to develop root rot tolerant cultivars. A complementary genetics and gene expression approach was undertaken in this study to identify *Fsp*-responsive genes in four tolerant and four susceptible pea genotypes. Time course RNAseq was performed on both sets of genotypes after *Fsp* challenge. Analysis of the transcriptome data resulted in the identification of 42,905 differentially expressed contigs (DECs). Interestingly, the vast majority of DECs were overexpressed in the susceptible genotypes at all sampling time points, rather than in the tolerant genotypes. Gene expression and GO enrichment analyses revealed genes coding for receptor-mediated endocytosis, sugar transporters, salicylic acid synthesis and signaling, and cell death were overexpressed in the susceptible genotypes. In the tolerant genotypes, genes involved in exocytosis, and secretion by cell, the anthocyanin synthesis pathway, as well as the DRR230 gene, a pathogenesis-related (PR) gene, were overexpressed. The complementary genetic and RNAseq approach has yielded a set of potential genes that could be targeted for improved tolerance against root rot in *P. sativum. Fsp* challenge produced a futile transcriptomic response in the susceptible genotypes. This type of response is hypothesized to be related to the speed at which the pathogen infestation advances in the susceptible genotypes, and the preexisting level of disease-preparedness in the tolerant genotypes.

## Introduction

The food industry contributes significantly to the world’s total greenhouse gas emissions (Poore and Nemecek, 2018). About 18% of the global greenhouse gas emissions are caused by livestock production, which supplies the majority of the dietary protein (Stehfest et al., 2009). Proposed mitigation efforts include a shift to plant-based protein as it is an environmentally sustainable option. The demand for plant-based protein is on the rise due to its health benefits (World Health Organization, 2015), as well as due to the ethical concerns related with exploiting animals as a source of protein (Johansson, 2019). The global plant-based protein market is expected to keep growing at a compound annual growth rate of 8.1% from 2019 to 2025 (Research and Markets, 2019). Popular plant-based meats from Impossible Foods and Beyond Meat have already reached some of the biggest food and retail brands in the US.

Pea (*Pisum sativum* L.; Family Fabaceae) is a major contributor to this plant-derived protein market and has gained particular interest lately due to its high content of lysine and tryptophan, overall high nutritional value and relatively low cost (do Carmo et al., 2016; Peng et al., 2016; Xiong et al., 2018). The market for pea protein is expected to be $34.8 million in 2020 due to growing consumer interest in plant-based proteins as an alternative to animal-derived protein (Grand View Research, 2015; Pietrysiak et al., 2018). Pea also plays a critical role in sustainable agriculture, due to its nitrogen-fixing capability, short life cycle, and low water usage; all of which make it a useful cover-and-break crop requiring minimal external inputs.

The US is one of the world’s major pea producers. In the US, harvested area of peas has increased by over 300% during the last 25 years; however, the yields have decreased an average of 7.5% throughout this timespan (Vandemark et al., 2014). Sustainable production of pea has been negatively affected by several diseases, predominantly root rots (Akhtar and Azam, 2014; Bodah et al., 2016). Root rots are the diseases of greatest impact to crop production worldwide (Kumari and Katoch, 2020). Frequently, root rot diseases involve more than one pathogen; therefore, the disease is known as root rot complex. Pathogens such as bacteria, oomycetes and fungi are commonly involved in this root rot complex (Xu et al., 2012; Chittem et al., 2015; Gossen et al., 2016).

One of the predominant causal agent of root rots in *P. sativum* is the soil fungus, *Fusarium solani* f. sp. *pisi* (*Fsp*). *F. solani* is a fungal soil-borne facultative parasite that is present worldwide (Zhang et al., 2006). The yields of *P. sativum* cultivars can be reduced by 15-60% by *Fsp* (Seaman, 1976; Grünwald et al., 2003; Porter et al., 2014). Over the years, hundreds of pea cultivars and germplasm core collections have been screened for *Fsp* resistance, and lines have been developed that demonstrate partial resistance to selected *Fsp* races (Coyne et al., 2008). An effort to identify tolerance to root rot in wild pea germplasm resulted in the identification of eight accessions with high levels of partial resistance (Porter, 2010). These accessions have been utilized for developing new cultivars. However, in tests replicated in the greenhouse and/or the field with derived selections, complete tolerance to *Fsp* has not been obtained (Grünwald et al., 2003; Porter et al., 2014; Bodah et al., 2016).

Understanding the genetic basis of tolerance to *Fsp* in a wide array of different pea breeding lines and cultivars has been pursued in several studies. The first QTL for *Fsp* tolerance was reported from a field study utilizing various parental lines that showed resistance to multiple root rots (Kraft, 1992; Feng et al., 2011). Recent studies conducted under controlled conditions have reported three QTLs; QTL *Fsp-Ps 2.1* explains 44.4 – 53.4% of the phenotypic variance within a 1.2 cM confidence interval. The other two QTLs, *Fsp-Ps 3.2* and *Fsp-Ps 3.3* explain 3.6-4.6% of the phenotypic variance related *Fsp* root rot tolerance (Coyne et al., 2015, 2019). While, the genes underlying these QTLs have not yet been identified, there is reason for optimism given the recent release of the pea reference genome (Kreplak et al., 2019). It is expected to facilitate characterization of potential transcription factors, stress-associated phytohormone genes, Pathogenesis-related (PR) proteins, or pea phytoalexin Pisatin (Kendra and Hadwiger, 1984) in the interaction between pea and *Fsp*.

While genetic approaches for identifying disease tolerance or resistance genes are common, gene expression approaches to identify key genes in response to pathogen challenge remain scarce. A report of *Aphanomyces euteiches*-mediated root rot of pea was investigated using a gene expression approach, and novel genes responsive during the pathogenic interaction with *Medicago truncatula* were reported (Nyamsuren et al., 2003). Besides the expected induction of pathogenesis-related (PR) and defense genes, several novel genes were also reported to be overexpressed during the plant-pathogen interaction.

To the best of our knowledge, a gene expression approach to identify genes involved in *Fsp* tolerance in pea is yet to be reported. For gaining a comprehensive insight into the transcriptomic responses during *Fsp* challenge, a comparative time-course RNAseq expression analysis was performed on four tolerant and four susceptible *P. sativum* genotypes that were selected from a preceding study (Bodah et al., 2016). Data analysis reaffirmed the role of Disease-Resistance Response 230 (DRR230) and sugar transporters, as well as expression patterns of genes associated with receptor-mediated endocytosis and exocytosis, cell death, and anthocyanin synthesis. Interestingly, several previously uncharacterized genes were also identified to be differentially expressed in both tolerant and susceptible genotypes, which may help illuminate novel mechanism of pea-*Fsp* interaction.

## Materials and methods

### Plant material and *Fsp* isolates

A total of eight, white-flowered pea genotypes were selected for pathogen challenge (Table 1). Four tolerant genotypes—00-5001, 00-5003, 00-5004, and 00-5007—were selected from the *Fsp* tolerant 5000 series (Porter et al., 2014). Four susceptible genotypes— ‘Aragorn’, ‘Banner’, ‘Bolero’, and ‘DSP’—were identified among frequently used commercial pea varieties. These eight genotypes were selected based on their root disease severity index reported in a preceding study (Bodah et al., 2016).

**Table 1:** Selected white-flowered pea genotypes for time course transcriptome analysis in response to *Fusarium solani* f. sp. *pisi* (*Fsp*) challenge. The table summarizes pea genotypes, source, Fsp tolerance level, other disease resistance, 100 seed weight, leaf type, and market class.

| Genotype | Source<sup>a</sup> | *Fsp* resistance level<sup>b</sup> | Other disease resistance<sup>c</sup> | 100 seed weight | Leaf type<sup>d</sup> | Market Class |
| --- | --- | --- | --- | --- | --- | --- |
| 00-5001 | USDA-ARS VFCRU | * | Fop races 1, 2 and 5 | 22.7 | af | Green fresh |
| 00-5003 | USDA-ARS VFCRU | * | Fop races 1, 2 and 5 | 15.9 | af | Green fresh |
| 00-5004 | USDA-ARS VFCRU | * | Fop races 1, 2 and 5 | 20.8 | af | Green fresh |
| 00-5007 | USDA-ARS VFCRU | * | Fop races 1, 2 and 5 | 22.2 | P | Green fresh |
| Aragorn' | ProGene | *** | Fop races 1, 2; PSBMV | 19.5 | af | Green dry |
| Banner' | ProGene | *** | Fop race 2, PM | 18.7 | af | Green dry |
| Bolero' | AsGrow | **** | Fop race 1, PM,<br>Pythium, EMV | 20.12 | P | Green fresh |
| DSP' | Canner Seed | *** | - | 20.9 | P | Green fresh |

<sup>a</sup>AsGrow = AsGrow Seed Co., San Juan Bautista, CA; Canner Seed=Canner Seed Co., Idaho Falls, ID; ProGene = ProGene LLC Plant Research, Othello, WA; USDA-ARS VFCRU=USDA-ARS, Vegetable and Forage Crops Research Unit, Prosser, WA.
<sup>b</sup>Fsp tolerance (Bodah et al. 2016)
<sup>c</sup>Fop= *Fusarium oxysporum*; PSBMV= Pea Seed-Borne Mosaic Virus, PM= Powdery Mildew; EMV= Enation Mosaic Virus
<sup>d</sup>af (Afila) = semi-leafless; P (Perfection) = normal leaf type

The *Fsp* isolates Fs 02, Fs 07, and Fs 09, were obtained from the Palouse Region of WA and ID, US soils by Dr. Lyndon Porter, USDA-ARS Vegetable and Forage Crops Research Unit, Prosser, WA. The three isolates were grown on pentachloronitrobenzene (PCNB) selective media for six days (Nash and Snyder, 1962). Cultures were transferred to KERR’s media (Kerr, 1963), and incubated on a shaker at 120 rpm under continuous light for six days at 23 to 25°C. The spore concentration of each isolate was determined using a hemocytometer and diluted to 1×10^6^ spores/ml of water. A spore suspension inoculum containing equal parts by volume of each of the three isolates was created.

### *Fsp* disease challenge

Seeds of each pea genotype were sterilized in a 0.6% sodium hypochlorite solution and rinsed in sterile distilled H2O. Seeds were then soaked for 16 hours in either the *Fsp* spore suspension (inoculated set) or in sterile H2O (control set). After the challenge with the spore suspension, seeds were harvested at specific time points of 0, 6, 12 hours (hr). The 0 hr time point began at the termination of the 16 hr inoculation period. Six hundred seeds were harvested per genotype per time point, immediately frozen under liquid nitrogen and transferred to storage at -80 °C for subsequent RNA extraction. The experiment was repeated three times.

### RNA isolation, cDNA library construction and sequencing

The frozen seed material was pulverized in a SPEX SamplePrep 6870 FreezerMill (SPEX SamplePrep, NJ, USA) for five cycles. Each cycle consisted of cooling for two minutes and grinding at 15 counts per second for four minutes. Total RNA was isolated from the pulverized tissue using the RNeasy Plant RNA Extraction Kit (Qiagen, Hilden, Germany). A Nanodrop ND-8000 Spectrophotometer (ThermoFisher, MA, USA) and a Qubit Fluorometer (Life Technologies, CA, USA) were used to quantify the extracted RNA. Contaminating DNA was removed using the TURBO DNA-free™ Kit (Life Technologies, CA, USA) using the manufacturer’s instructions. RNA quality was verified via electrophoresis on a 1% agarose gel.

Equimolar amounts of RNA samples from tolerant and susceptible genotypes were bulked for each time point prior to the construction of RNAseq libraries. RNAseq libraries were constructed using 1 μg of RNA, and the Illumina TruSeq kits (Illumina Inc. San Diego, CA, USA). RNA was purified with an Oligo(dT) cellulose affinity matrix, and subsequently fragmented into short pieces of an average size of 450 base pairs with Ampure XP beads (Beckman Coulter, CA, USA). All libraries were quantified on a Qubit Fluorometer (Life Technologies, CA, USA) and analyzed on an Agilent BioAnalyzer (Agilent Technologies, CA, USA) to determine concentration, final size and purity of the library. A total of 24 libraries were sequenced using the HiSeq2000 configuration 100 PE (Illumina Inc. CA, USA) at the Michigan State University Genomics core laboratory.

### RNAseq data processing and statistical analysis

The generated fastq files were analyzed for quality with CLC Bio Genomics Workbench 6.0.1(CLC Bio, Aarhus, Denmark) and trimmed with trimmomatic (Bolger et al., 2014). *De novo* RNAseq assembly was performed using data from all 24 samples to obtain a master assembly with the software Trinity v2.8.4 (Grabherr et al., 2011). Dependencies for Trinity, Bowtie2 v1.2.3 (Langmead and Salzberg, 2012), Salmon v0.12.0 (Patro et al., 2017), and JELLYFISH v2.2.3 (Marçais and Kingsford, 2011), were used during assembly. Bowtie2 and Salmon were used for abundance estimation, and JELLYFISH was used as a k-mer counting software.

The software Kalisto was used for transcript quantification (Bray et al., 2016). The reads were quantified for each of the two biological replicates of the tolerant or susceptible genotypes at three time points, 0, 6, 12 hr. after inoculation, and for each control or treatment. This analysis resulted in 24 separate quantification groups that were used for comparison. Using Baggerley’s test, differentially expressed contig (DECs) with p value of <0.001 and a greater than 2-fold change in expression were identified. The RPKM (Reads Per Kilobase of transcript per Million mapped reads) expression values were also ascertained for each contig. Heat maps showing fold-change of RPKM values between control (C) and inoculated (I) sets and among genotypes were created in Microsoft Excel 365 ProPlus (Microsoft Corporation, WA, US).

### Functional annotation, Statistical GO enrichment and pathway analysis

Functional annotation of the master assembly and DECs was conducted via BLAST in BLAST2GO v. 3.3. (Conesa and Götz, 2008). Default parameters were used for the functional annotation, as well as for GO mapping, and InterPro Scan. The two-tailed Fisher’s exact test (FDR < 0.05) was used to ascertain over- and under-represented functions during *Fsp* challenge. A heat map representing ‘biological process’ GO terms over-represented in the *Fsp* inoculated treatment was created in Microsoft Excel 365 ProPlus (Microsoft Corporation, WA, US).The Kyoto Encyclopedia of Genes and Genomes (KEGG) pathway analysis was performed to identify pathways represented by the set of DECs for each time point and genotype.

### Real-time quantitative PCR

RNA was extracted utilizing the RNeasy Plant DNA Extraction Kit (Qiagen, Mainz, Germany) from the same sampled tissues utilized for the RNAseq analysis. After DNase treatment, equimolar amounts of RNA from the tolerant and susceptible genotypes were bulked for each time point. First-strand cDNA synthesis was performed using 1,500 ng of each bulked RNA sample with the SuperScript ® Vilo kit (ThermoFisher Scientific, MA, US). Nine genes were randomly selected from the list of DECs for RT-qPCR analysis (Table S1). Primers for RT-qPCR were designed with the Primer3 software (Rozen and Skaletsky, 2000) with the corresponding transcriptome contig as the query sequence for each primer set. The *Pisum sativum* root border cell specific protein (GenBank accession AF1139187.1) was used as an internal reference control as it showed invariant expression across genotypes and treatments in the RNAseq data.

The QUBIT 3.0 fluorometer (Invitrogen, CA, US) was used to quantify cDNA library concentration. For each reaction, 16ng of cDNA was used with the iTaq™ Universal SYBR® Green Supermix (BIO-RAD, CA, US). Each RT-qPCR reaction was performed in triplicate for each of the three biological replicates using the Stratagene Mx3005P (ThermoFisher Scientific, MA, US). The amplification profile consisted of an initial denaturation at 95 °C for 150 seconds, 40 cycles of 20 seconds at 95 °C for denaturation, 20 seconds at 60 °C for annealing, and 20 seconds at 72 °C for extension. A melting curve analysis was performed post amplification to ensure the presence of a unique amplicon and performed with an initial denaturation at 95 °C for 1 min and a decrease of temperature to 50°C for annealing. Temperature was then increased in 0.5°C increments at 5 sec/step from 50 °C to 95 °C for fluorescence readings. Raw fluorescence data was used as input for crossover threshold (Ct) calculations and reaction efficiencies adjusted with LinRegPCR 2012.0 software (Ruijter et al., 2009). The ΔΔCt method offered by PE Applied Biosystems (Perkin Elmer, Forster City, CA) was used to obtain relative differential expression values after reaction efficiencies were adjusted with the LinRegPCR 2012.0 software (Pfaffl, 2001).

### Functional annotation of QTL associated with *Fsp* tolerance in pea

*Fsp-Ps 2.1*, the major QTL found to be associated with *Fsp* tolerance in pea (Coyne et al., 2019), was annotated using the transcriptome data generated in this study to determine if there are any differentially expressed genes located in the selected genomic region. *Fsp-Ps 2.1* explains 44.4 – 53.4% of the phenotypic variance and it is located on chromosome II within a 1.2 cM confidence interval of maker Ps900203 (Duarte et al., 2014; Coyne et al., 2019). The genomic sequence of the *Fsp-Ps2.1* ± 1.2 cM (±201,800 nt) was obtained from the pea genome (Kreplak et al., 2019). The length of the QTL sequence was calculated based on the distance between the Ps900203 and Ps000075 markers. Marker Ps000075 is 0.6 cM=168,166 nt away from Ps900203. The transcriptome data from this study, was aligned via BLAST against the *Fsp-Ps2.1* ± 1.2 cM sequence in CLC Bio Genomics Workbench 6.0.1 (CLC Bio, Aarhus, Denmark).

## Results and Discussion

### Assembly of transcriptome data and identification of Differentially Expressed Contigs (DECs)

A total of 850 million reads were generated after sequencing of the 24 libraries (Table S2). For these 24 libraries, the mean Q score ranged from 34.00 to 34.88; these Q scores validate the quality of the assay (Table S2). After QC and trimming of low-quality reads, 69.49% of reads were used for assembly of the master transcriptome. The master transcriptome, generated in this study, was composed of 185,721 contigs (Table S3), which had a mean contig length of 1503.15 nucleotides (nt) with a length range of 184–18,990 nt.

Mapping of reads to the master assembly showed a different number of contigs with zero mapped reads for each time point (0, 6, 12 hr.) and genotype (tolerant and susceptible). After expunging contigs with zero mapped reads, the total number of contigs ranged from 102,382 to 141,530, or 55.13 to 76.21% of the total 185,721 contigs, respectively (Table S2). The total number of reads mapped to each contig for each time point and genotype, and RPKM values are summarized in Table S3.

For each contig, twelve pairwise comparisons were performed (Table S4 and Table S3). In order to identify which genes were differentially expressed in response to the *Fsp* challenge, six comparisons, as listed in Table S4-comparisons 1 to 6, were performed (Fig 1a). In order to identify which genes responded differentially to *Fsp* between the tolerant and susceptible genotypes, six additional pairwise comparisons were performed and are summarized in Table S4-comparisons 7 to 12 (Fig 1b). The twelve pairwise comparisons resulted in the identification of 42,905 DECs out of the 185,721 contigs.

**Figure 1.**
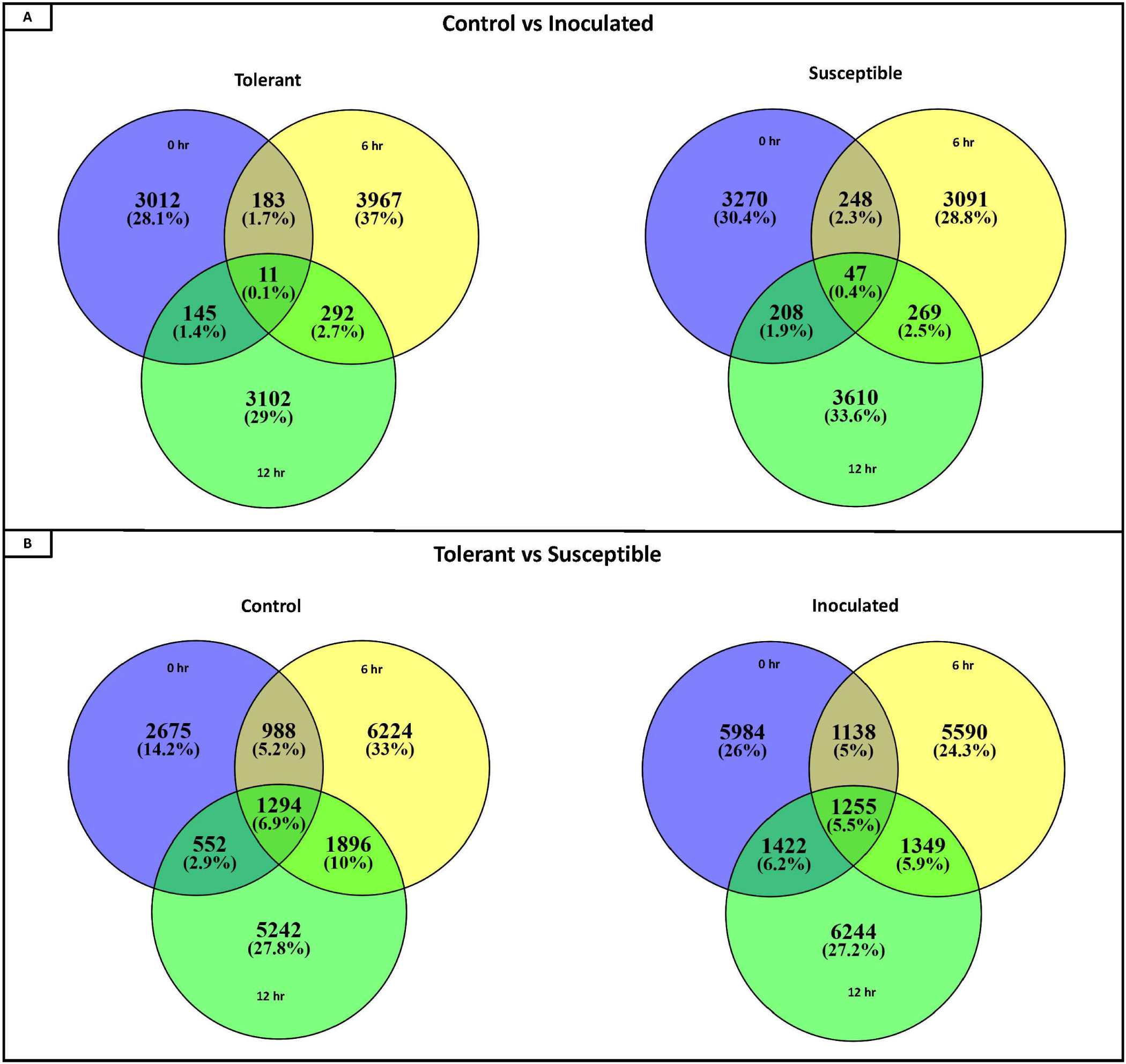
Venn diagrams representing number of DECs (Differentially Expressed Contigs) for the twelve pairwise comparisons. (a) Number of DECs for pairwise comparisons between control and inoculated samples collected at 0, 6 and 12 hr time points for the tolerant and the susceptible genotypes. (b) Number of DECs for pairwise comparisons between the tolerant and susceptible genotypes for each time point (0, 6 and 12 hr) for control and inoculated conditions.

Pairwise comparisons 1 to 6, yielded the number of upregulated DECs obtained for each time point (Fig 2). For the *Fsp* inoculated tolerant genotypes, the number of upregulated DECs varied between 1,200 and 1,460 DECs across 0, 6 and 12 hr time points. The number of suppressed (under-expressed) DECs in the inoculated tolerant genotypes was larger (2795-4453 DECs). For the *Fsp* inoculated susceptible genotypes, the total number of upregulated DECs was 5-7 times larger than the total number for the *Fsp* inoculated tolerant genotypes (Fig 2). The number of suppressed genes in both sets of genotypes was similar at 0 hr. However, for the susceptible genotypes, these numbers were 1.34 and 2.21 times larger than the tolerant genotypes at 6 and 12 hr, respectively. A total of 5.7 and 5.4% of the DECs were shared across the three time points for the tolerant and susceptible genotypes, respectively.

**Figure 2.**
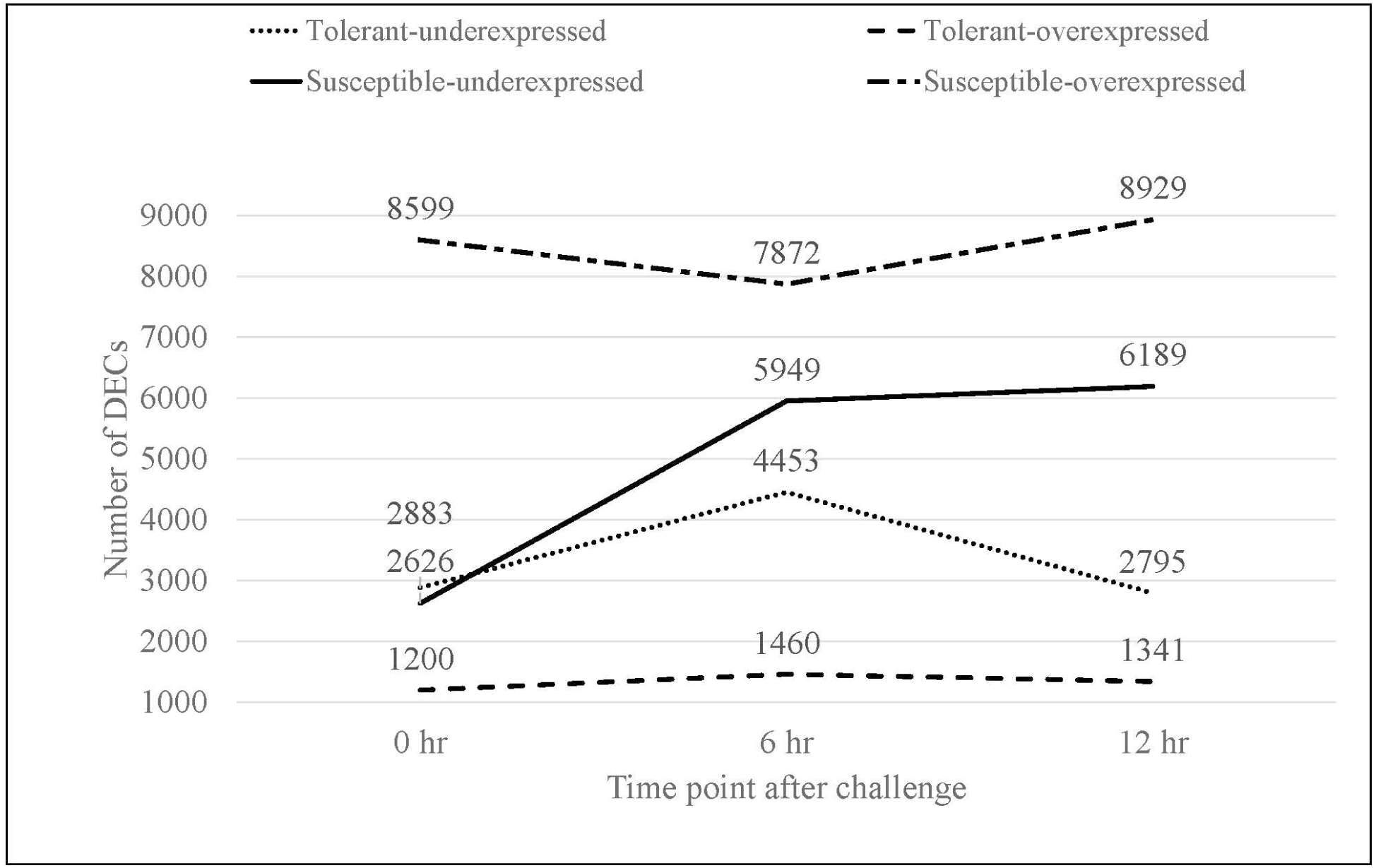
Total number of overexpressed and underexpressed DECs in the inoculated treatments for tolerant and susceptible genotypes at each time point in response to *Fsp* challenge.

The larger number of upregulated DECs represented a more diverse and higher number of transcriptionally regulated genes in the susceptible genotypes, in contrast to the tolerant genotypes. This higher number of upregulated DECs in the susceptible genotypes may be due to involvement of various biological processes associated with successful *Fsp* infection in pea embryonic tissue. This observation is consistent with previous studies, which showed that pathogen attack engages a broader range of pathways and a larger proportion of genes in the susceptible genotypes compared to the resistant ones (Bagnaresi et al., 2012; Zheng et al., 2013; Matic et al., 2016).

From the total upregulated DECs in the *Fsp* inoculated sets, only 10 (0.10%), 48 (0.52%), and 12 (0.12%) DECs are shared among the tolerant and susceptible genotypes at the 0, 6, and 12 hr time points, respectively. Therefore, the response mechanisms involved in the *Fsp* challenge were divergent between the tolerant and susceptible genotypes. The large difference in overexpressed DECs could explain the difference in tolerance between these sets of genotypes. The genes that were overexpressed in susceptible genotypes were numerically different from the ones in the tolerant genotypes.

### Functional annotation of DECs and GO-term Enrichment

From the 185,721 contigs of the master assembly, 120,132 returned positive BLAST hits when queried to the NCBI database (National Center for Biotechnology Information). A total of 4,734 contigs, or 2.55% of the total contigs were annotated as proteins of unknown function or hypothetical proteins. The top BLAST hits showed similarity to *Medicago truncatula, Trifolium pratense, Cicer arietinum*, and *Trifolium subterraneum* with a distribution of 34.0%, 16.1%, 16.0, and 14.6% respectively. *Pisum sativum*, with a 3.6% match, was fifth in the rankings. The low percentage of hits to *P. sativum* was most likely due to the relatively scarce number, in comparison to the crops mentioned previously, of transcriptomic studies of *P. sativum* represented in the NCBI database. Of the 42,905 DECs, 36,923 (86.1%) returned positive BLAST hits when aligned to the NCBI database. Interestingly, 33 contigs (0.09%) of the 36,923 hits were classified as proteins of unknown function or hypothetical proteins, which could be useful candidates for understanding the pea-*Fsp* interaction.

The GO enrichment analysis identified significant over- and under-represented GO terms for each of the three *Fsp* inoculated time points (Table S5). GO terms related to routine DNA processes, such as DNA metabolic process, DNA biosynthetic process, and DNA integration, were significantly underrepresented at different time points in the tolerant and susceptible genotypes. DNA integration and DNA metabolic process terms were underrepresented across the three times in the susceptible genotypes, but only at 0 and 6 hr for the tolerant genotypes. Nucleic acid phosphodiester bond hydrolysis was also underrepresented only during certain times points in the susceptible genotypes but not in the tolerant genotypes.

Figure 3 represents a heat map with overrepresented biological process GO terms for each time point in the inoculated treatment for the tolerant and susceptible genotypes. These terms provide a comparative perspective of biological processes that responded to *Fsp* in the two subsets of genotypes. Common terms related to basic plant metabolism, such as gene expression, regulation of primary metabolism, transcription, and protein synthesis were over-represented at the 0 hr time point in the tolerant genotypes. Conversely, these terms were upregulated in at least 2 time points in the susceptible genotypes. Therefore, these results showed the tolerant genotypes may have moved towards a basal metabolic state after the 0 hr time point while the susceptible genotypes were responding to *Fsp* throughout the entire time course presented in this study.

**Figure 3.**
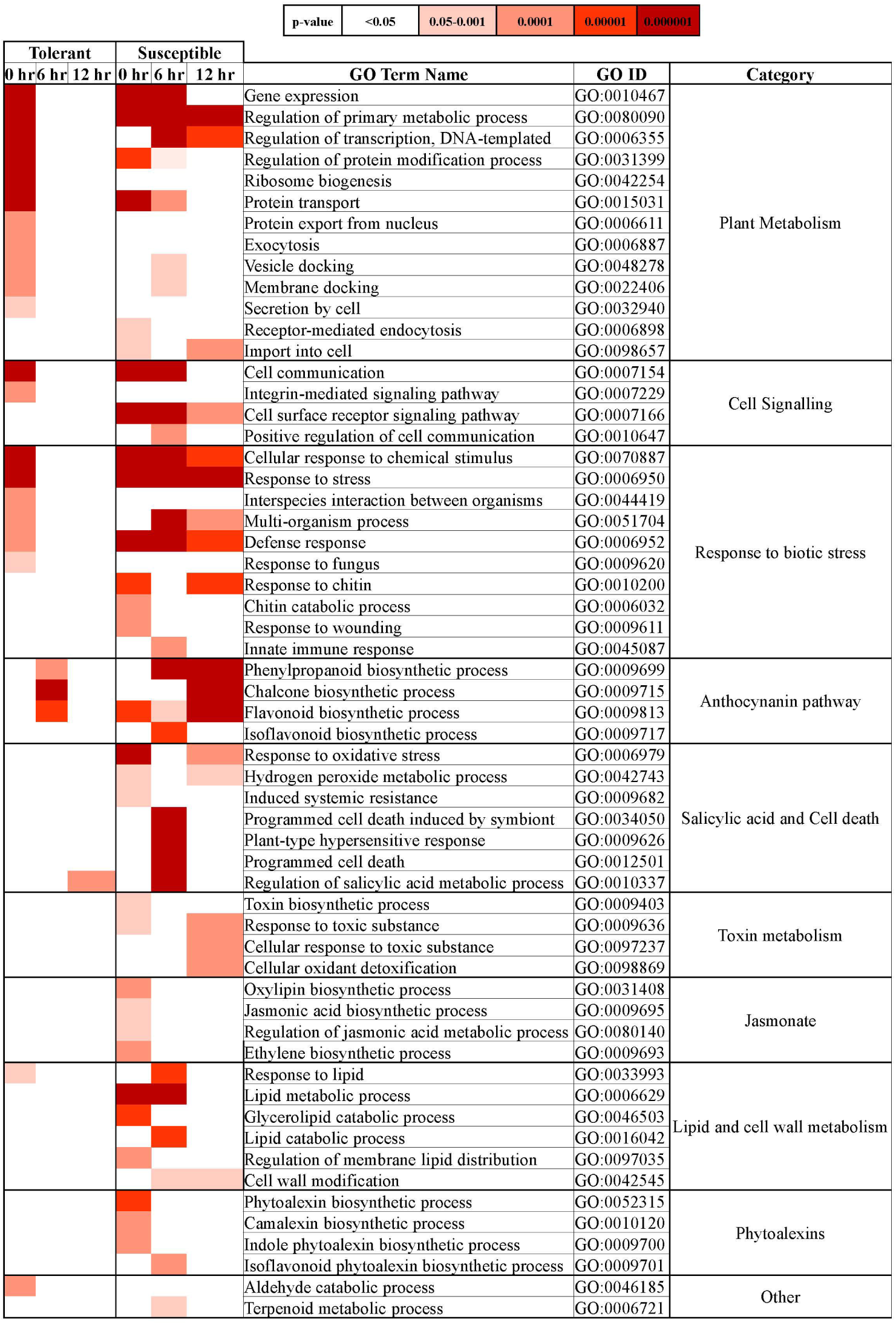
Biological process-GO terms over-represented in the *Fsp* inoculated treatment for a tolerant and susceptible genotypes at 0, 6, and 12 hr time points. The tone of colors in the heatmap denotes the p-value of the fold-change in expression, as indicated in the key. Significantly over-represented GO terms showed a p>0.05.

GO terms such as ribosome biogenesis, protein export from nucleus, exocytosis, and secretion by cell were over-represented at 0 hr in the tolerant genotypes. Plants are known to transport antimicrobial molecules, such as peptides and/or secondary metabolites, outside the cell to function in plant immunity (Kwon and Yun, 2014). In contrast, the susceptible genotypes seemed to be importing substances inside the cell since terms such as receptor-mediated endocytosis and import to cell were over-represented. Endocytosis seems not only to play a role in pathogen-associated molecular patterns (PAMP)-triggered immunity and effector-triggered immunity but also in susceptibility. Vesicle endocytosis can be manipulated by pathogens and can import pathogen-derived effectors into the plant cell (Driouich et al., 1997; Kwon and Yun, 2014). In this study, while the tolerant genotypes seemed to export antimicrobial molecules potentially to counter *Fsp*, the susceptible genotypes seemed to import substances. Thus, it is hypothesized that the suppression of exocytosis mechanisms in the susceptible genotypes might block or delay the transport and release of antimicrobial substances against pathogens.

GO terms in the cell signaling and response to biotic stress categories also showed an early and unique response at 0 hr in the tolerant genotypes, while this response was present throughout the entire experiment (0, 6 and 12 hr) or at later stages (6 or 12 hr) in susceptible genotypes. Several studies have analyzed how pea responds to *Fsp* and the non-host pathogen *F. solani* f. sp. *phaseoli* (*Fsph*). These studies concluded that the major difference is the speed at which the pea plants react. The type of response exhibited by pea varies with the rate of induction of PR genes and other associated biochemical pathways. In the case of either *Fsph* or *Fsp* infection, the fungus releases DNAses, which localize to the host nuclei and digest the nuclear DNA (Hadwiger and Adams, 1978; Hadwiger, 2008, 2015). Fungal DNases can also impact the nuclei in the fungal mycelia and trigger their deterioration (Hadwiger, 2008). In the case of a compatible interaction (successful infection leading to disease) between *Fsp* and pea, the slower reaction rate of the pea host allows *Fsp* to protect a small number of its own nuclei from fungal DNAses. The slower reaction allows the growth of *Fsp* to resume after 12 hours post-inoculation (Klosterman et al., 2001; Hadwiger, 2015). In contrast, the relatively rapid response generated in the host against *Fsph* terminates the growth of the fungi at 6 hrs. post-inoculation (Hadwiger, 2008, 2015). Given this information, it is hypothesized the speed of reaction to the pathogen may be one of the mechanisms of tolerance in the tolerant genotypes.

The GO terms associated with salicylic acid and cell death category were, in most cases, overrepresented only in the susceptible genotypes. Therefore, the susceptible genotypes were expected to have a more intense response to *Fsp* through induced systemic resistance, host programmed cell death, and plant-type hypersensitive response.

The GO terms associated with the production of flavanones, flavones, flavonols, proanthocyanidins, and anthocyanins showed over-representation in both genotypes. In the tolerant genotypes, these terms were only over-represented at 6 hrs. In the susceptible genotypes, GO terms from this category were overrepresented at the 0, 6 and 12 hr time points, indicating a more intense response. GO terms associated with the jasmonate pathway, phytoalexin synthesis, toxin metabolism and lipid metabolism were overrepresented only in the susceptible genotypes at all timepoints (Figure 3 and Table S5).

### Quantitative RT-PCR Verification

To verify the expression results obtained from RNAseq data, RT-qPCR analysis was performed on nine randomly selected genes at all time points for the tolerant and susceptible genotypes. The expression trends of eight out of nine genes (89%) correlated with the RPKM values indicating the robustness of RNAseq results (Fig 4). The internal reference control (GenBank accession AF1139187.1) showed invariant expression across genotypes and treatments in this RT-qPCR analysis.

**Figure 4.**
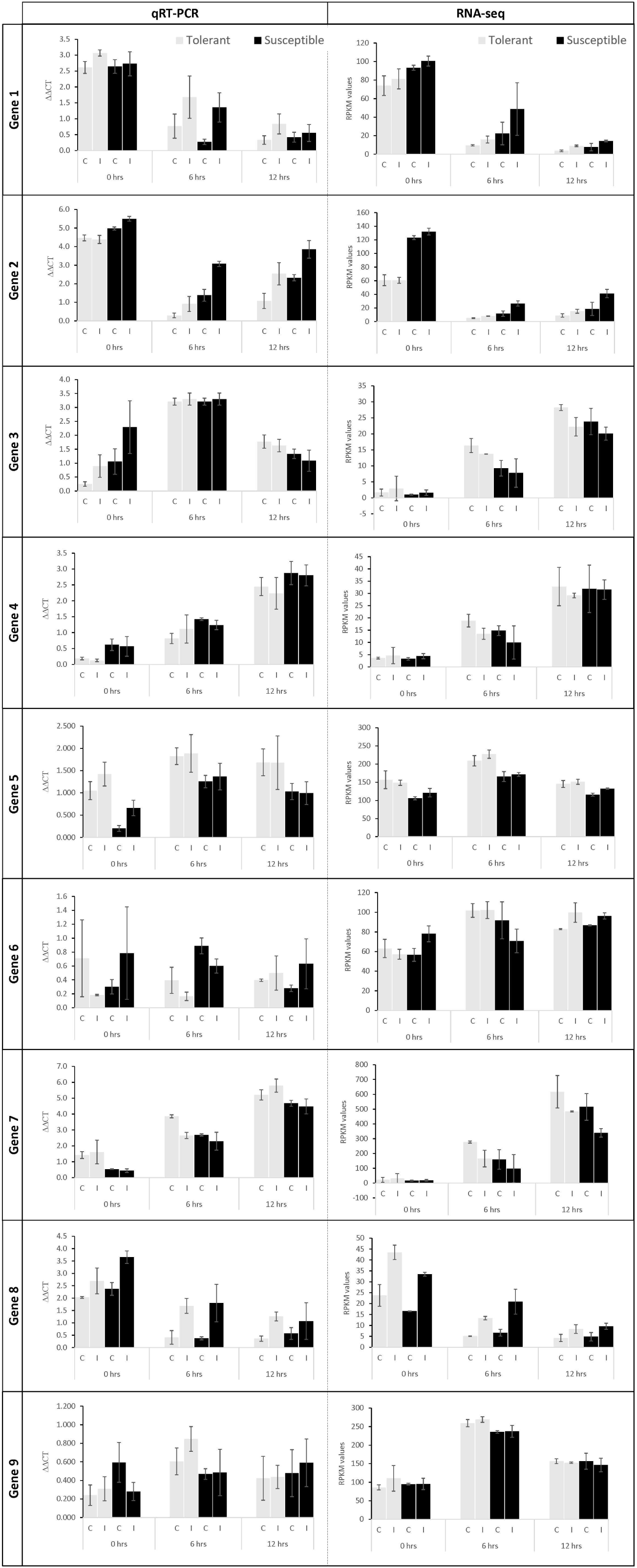
RT-qPCR validation of select genes in control (C) and Fsp inoculated (I) plants. The grey and black bars represent relative gene expression for the tolerant and susceptible genotypes, respectively. First column: RT-qPCR data show the average relative expression of three biological samples with three technical replicates each. Second column: RPKM values calculated for each gene. The error bars represent the standard error between replicates in RT- qPCR analysis. Gene 1: TRINITY_DN2419_c0_g1_i5 (BLAST accession: XM_003592027.3), Gene 2: TRINITY_DN2754_c0_g1_i11 (BLAST accession: XM_004502933.3), Gene 3: TRINITY_DN5727_c0_g1_i1 (BLAST accession: MK618561.1), Gene 4: TRINITY_DN6240_c0_g1_i9 (BLAST accession: XM_003592048.3), Gene 5: TRINITY_DN2169_c1_g1_i2 (BLAST accession: XM_004506541.3), Gene 6: TRINITY_DN1232_c0_g1_i11 (BLAST accession: XM_004504351.3), Gene 7: TRINITY_DN5529_c0_g1_i9 (BLAST accession: XM_024782286.1), Gene 8: TRINITY_DN8631_c0_g1_i1 (BLAST accession: XM_013611166.2), Gene 9: TRINITY_DN1795_c0_g1_i2 (BLAST accession: XM_004514502.3).

### *Fsp*-induced differential gene expression

When a pathogen begins an interaction with a plant, several interconnected signaling and defense pathways are activated. The molecular response to Fusarium–plant interaction is diverse, with variations in interactions depending upon the specific host genotype involved in the process. A closer look at the 42,905 DECs showed that most of these genes are known to participate in defense responses and might play a significant role against the *Fsp* pathogenicity in pea. These genes were placed into seven broad categories: 1. Expression of signaling-related genes, 2.

Genes involved in transcriptional regulation, 3. Pathogenesis-related (PR) genes, 4. Anthocyanin and lignin biosynthetic pathway genes, 5. Sugar metabolism, 6. Phytohormones, 7. Cell wall and membrane metabolism, and toxin metabolism.

#### 1. Expression of signaling-related genes

Plants detect pathogens via host sensors known as pattern-recognition receptors (PRR), which act by detecting PAMPs. PAMPs are molecules shared by groups of related pathogens. These molecules are characteristic of a pathogen and are essential for the survival of those organisms, and are not found associated with plant cells (Moffett et al., 2002; Beck et al., 2012; Ao et al., 2014). After the detection of PAMPs, PRRs induce PAMP-triggered immunity (PTI).

To date, the identified PRRs are mainly single pass transmembrane proteins that carry leucine-rich repeats (LRRs) or a lysin motif (LysM). Both types of PRRs are receptor-like kinases (RLKs), which carry a cytosolic kinase domain; or receptor-like proteins (RLPs), which have a short cytoplasmic tail without a kinase domain (Beck et al., 2012). Beck et al., (2012) identified three PRRs that have been proven to be specific to fungi: the Chitin Elicitor Binding Protein (CEBiP), the chitin elicitor receptor kinase I (CERK1), and the ethylene-inducing xylanase (Eix2). CEBiP and CERK1 cooperatively regulate chitin elicitor signaling to activate plant defense system (Hayafune et al., 2014). Interestingly, CEBiP was not identified in this study. Contigs corresponding to CERK1 and Eix2 genes were identified, however they were not differentially expressed in any of the genotypes.

In this study, several LRR-RLK receptors were found to be differentially expressed in response to *Fsp* challenge (Table S3). Most of these genes were upregulated in the susceptible rather than in the tolerant genotypes. The L-type lectin-domain containing receptor kinase, proline-rich receptor-like protein kinase, cysteine-rich receptor-like protein kinase and the wall-associated receptor kinases are also upregulated in the susceptible genotypes, following the same trend as above. The activity of Mitogen-activated protein kinases (MAPK), MAPK kinases (MAPKK), and MAPKK kinases (MAPKKK) were, in most cases, upregulated in the tolerant and susceptible genotypes after the challenge with *Fsp* when compared to the control. However, the expression was, in most cases, significantly higher in the susceptible genotypes throughout the time-course when compared to the tolerant genotypes.

Two contigs identified as receptors were found to change significantly in expression after the *Fsp* challenge in the susceptible genotypes only. Contig DN1290_c0_g1_i9 and Contig DN7023_c0_g2_i5 were identified as a receptor-like cytoplasmic kinase 176 and CC-NBS-LRR resistance protein (Table S3), respectively. The receptor-like cytoplasmic kinase 176 acts downstream of the CERK1 gene in the fungal chitin signaling pathways that mediates innate immunity responses such as reactive oxygen species generation, defense gene expression, and callose deposition (Ao et al., 2014). The CC–NBS–LRR proteins initiate a resistance response that often includes a type of cell death known as the hypersensitive response (HR) (Moffett et al., 2002). In the susceptible genotypes, contig DN1290_c0_g1_i9 was significantly overexpressed at 6 hr (FC=3.19) after the *Fsp* challenge. Expression of Contig DN7023_c0_g2_i5 was found to be lowered at 12 hr (FC=-3.59) after the *Fsp* challenge. Interestingly, no change was observed in the expression of the two contigs in the tolerant genotypes. These two contigs also showed higher expression in the susceptible genotypes when compared to their expression in the tolerant genotypes. Contigs DN1290_c0_g1_i9 and DN7023_c0_g2_i5 were significantly upregulated at 0 (FC=3.49), 6 (FC=5.08), and 12 hr (FC=6.77), and 6 hr (FC=71.69), respectively, in the inoculated treatments of the susceptible genotypes when their expression values were compared to the tolerant genotypes. The data on the observed induction of genes coding for receptor-like cytoplasmic kinase 176 and CC-NBS-LRR resistance protein in the susceptible genotypes is intriguing. It suggests that the pathogen likely recruits oxygen species generation, hypersensitive response, defense gene expression, and callose deposition to establish infection. It raises a question if loss of function mutation in these genes in the susceptible genotypes could confer tolerance to *Fsp*. A loss-of-function mutation in receptor-like cytoplasmic kinases and CC-NBS-LRRs genes has proven to confer tolerance to different pathogens (Lorang et al., 2007; Sweat and Wolpert, 2007; Zhang et al., 2019).

#### 2. Genes involved in transcriptional regulation

Activation of the expression of disease-related transcription factors (TFs) plays a crucial role in disease resistance or susceptibility against pathogens (Berrocal-Lobo et al., 2002; Eulgem and Somssich, 2007). The following TFs were found to be differentially expressed between the tolerant and susceptible genotypes and/ or were influenced by *Fsp* challenge: bZIP, ERF, MYB, GATA, MADS-box, NAC, PLATZ, KAN2, PosF21, WRKY, C2H2, bHLH, DIVARICATA, E2F, GLABRA, ICE, IIIB, Jumonji, PIF, RF2a, SRM1, TCP19, TGA, UNE12, and HMG. From this list of TFs, bZIP, ERF, MYB, MADS-box, NAC, WRKY, C2H2, bHLH, E2F, Jumonji, PIF, RF2a, TCP19, TGA, and HMG have been reported to be master regulators of defense responses against pathogens (Pontier et al., 2001; Vailleau et al., 2002; Dong et al., 2003; Pré et al., 2008; Wang et al., 2009; Isaac et al., 2009; Kielbowicz-Matuk, 2012; Alves et al., 2013; Song et al., 2013; Li et al., 2013; Chandran et al., 2014; Khong et al., 2015; Li, 2015; Paik et al., 2017; Im et al., 2019). However, the involvement of following TFs in response to pathogen challenge has not been reported previously – GATA, PLATZ, KAN2, PosF21, DIVARICATA, GLABRA, ICE, IIIB, SRM1, UNE1.

In the tolerant genotypes, TFs were either not differentially expressed when the control and inoculated samples were compared, or their expression was significantly suppressed after *Fsp* challenge. In the susceptible genotypes, however, the expression of TFs remained the same or increased after *Fsp* challenge. When the inoculated treatments in the tolerant and the susceptible genotypes were compared at either time point, TFs were overexpressed in the susceptible genotypes (Figure 5).

**Figure 5.**
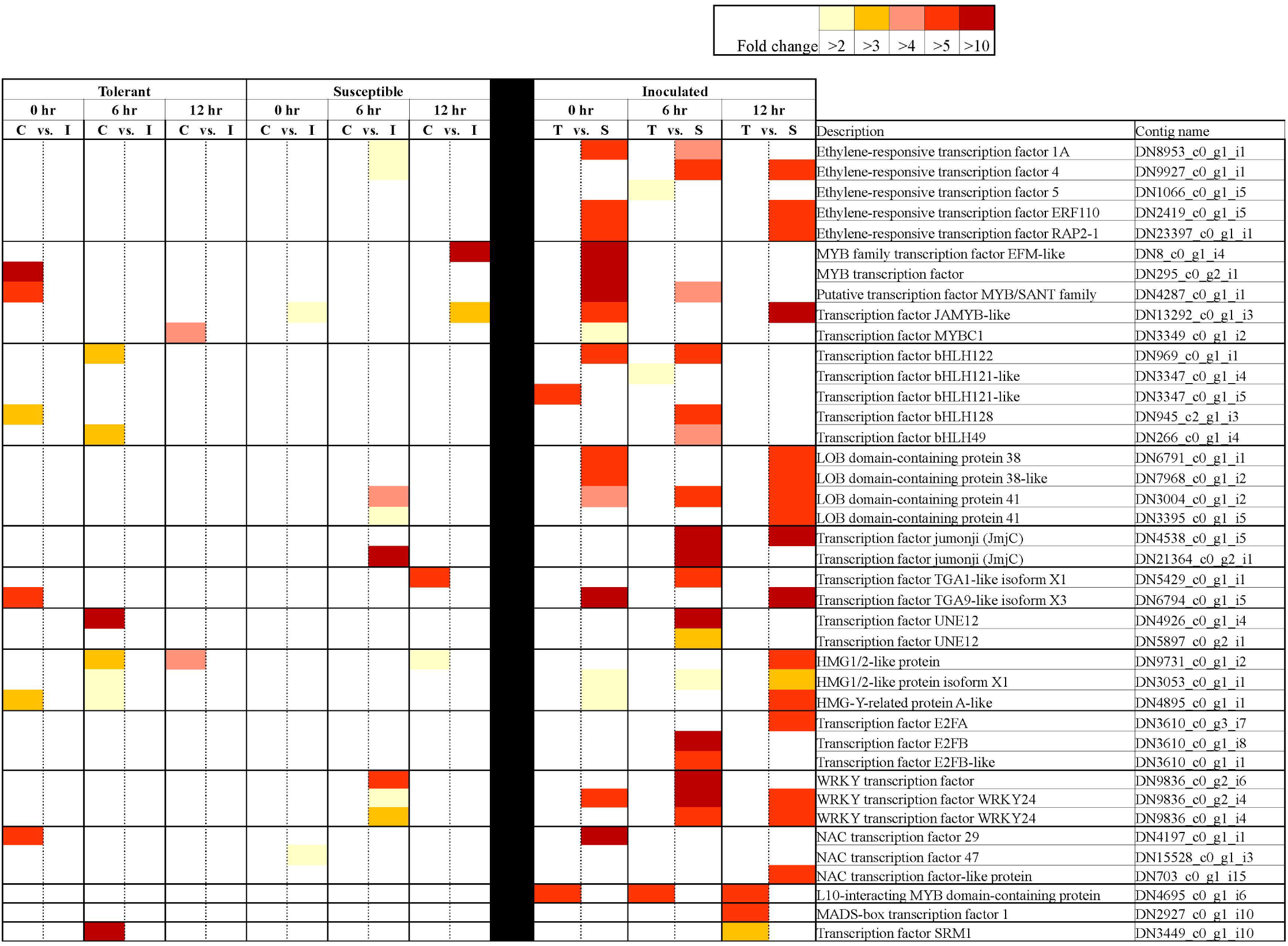
Differentially expressed transcription factors in tolerant and susceptible pea genotypes in response to *Fsp* challenge. The color key denotes fold-change. Pairwise comparisons that displayed greater than 2-fold difference (p> 0.005) in expression were identified with a color that ranges from light yellow (fold change>2) to dark red (fold change> 10).

The higher expression of TFs in the susceptible genotypes implies their role in susceptibility to *Fsp*. Indeed, it is well documented that overexpression of certain TFs causes susceptibility to certain pathogens (Kim et al., 2006; Lai et al., 2008; Thatcher et al., 2012). Some examples include the enhanced susceptibility of the *WRKY7*-overexpressing Arabidopsis plants to *Pseudomonas syringae* infection. Overexpression of *WRKY7* results in reduced expression of defense-related genes and a higher accumulation of salicylic acid (Kim et al., 2006). Similarly, overexpression of the AtWRKY4 gene in Arabidopsis enhances susceptibility towards the biotrophic bacterium *P. syringae* (Lai et al., 2008). In a disease screen with *F. oxysporum*, it was found that disruption of the *LATERAL ORGAN BOUNDARIES* (*LOB*) *DOMAIN* (*LBD*) TF led to increased resistance to *F. oxysporum* root-rot disease in *A. thaliana*. It has been suggested that Gene LBD20 functions in the jasmonate signaling pathway (Thatcher et al., 2012).

From the extensive list of differentially expressed TFs identified in this study, only the HMG A has been previously shown to participate during pea-*Fsp* interaction (Hadwiger, 2008). HMG A has been shown to complex with AT-rich regions within the promoter areas of Disease-Resistance Response (DRR) genes in pea after *Fsp* infection (Klosterman et al., 2003). Through interactions with the promoter area, HMG-I/Y positively or negatively regulates gene expression. In pea pod endocarp tissue, HMG-I/Y expression was observed at high levels in untreated tissue and at lower levels at 6 hr following *Fsp* inoculation or wounding of the tissue (Klosterman et al., 2003). Western blots also revealed that pea HMG-I/Y is expressed at decreased levels 6 h following *Fsp* challenge. Transient expression experiments also implicate the HMG-I/Y abundance in the down-regulation of DRR206 gene expression in pea (Klosterman et al., 2003). In this study, contigs DN4895 and DN12030 were identified as HMG-I/Y genes. Interestingly, the expression of the DN4895 contig decreases significantly at 0hr (FC=-3.59) and 6 hr (FC=-2.65) after *Fsp* challenge in the tolerant genotypes. On the other hand, the expression of this contig increased significantly at 6 hr (FC=3.51) after *Fsp* challenge in the susceptible genotypes. Furthermore, the expression of contig DN4895 was significantly higher under inoculated conditions at 0 hr (FC=2.56) and 12 hr (FC=7.91) for the susceptible genotypes compared to these expression values in the tolerant genotypes. Contig DN12030 is also significantly overexpressed under inoculated conditions in the susceptible genotypes at 0 hr (FC=3.23) and 12 hr (FC=5.70). It is plausible that HMG-I/Y mediates the reduction of expression of defense-related genes in the susceptible genotypes; however, this assumption will need to be further evaluated.

#### 3. Pathogenesis-related (PR) Genes

Pathogenesis-related (PR) proteins are expressed at basal levels in healthy plant tissues. Accumulation of PR proteins occurs only during a plant-pathogen interaction (Sels et al., 2008; Saboki Ebrahim and Singh, 2011). Plant PR proteins are low molecular weight and protease-resistant proteins, and have been categorized into 17 families according to their properties and functions such as glucanases, chitinases, ribosome-inactivating proteins, and defensins (Sels et al., 2008; Saboki Ebrahim and Singh, 2011). The inoculation of the tolerant and susceptible genotypes with *Fsp* generated changes in the expression of PR genes. Figure 6 shows DEC-PR genes identified in this study with their respective functions.

**Figure 6.**
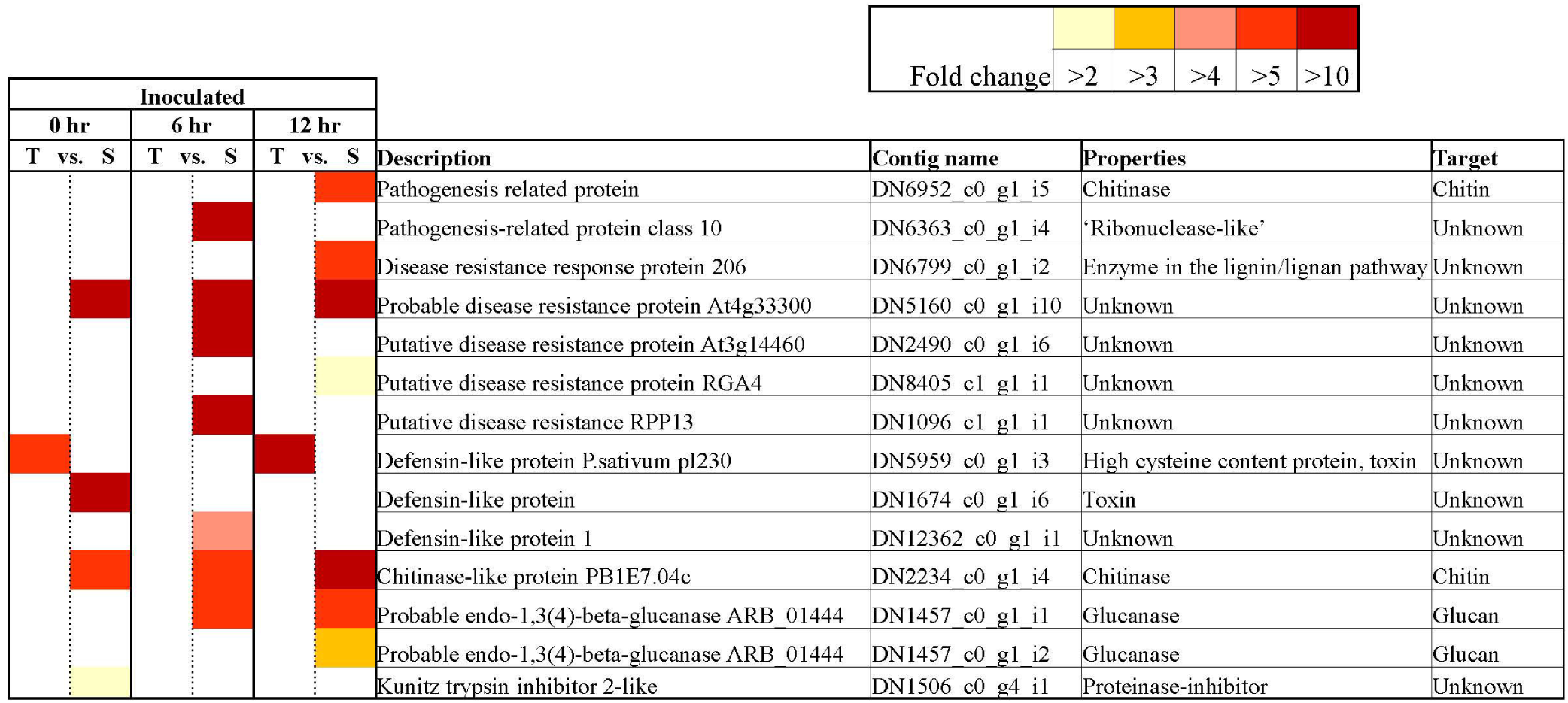
Key differentially expressed Pathogenesis-related contigs in tolerant and susceptible pea genotypes in response to *Fsp* challenge. The color key denotes fold-change. Pairwise comparisons that displayed greater than 2-fold difference (p> 0.005) in expression were identified with a color that ranges from light yellow (fold change>2) to dark red (fold change> 10).

All the PR protein encoding genes that were identified in this experiment were overexpressed in the susceptible genotypes over the tolerant genotypes except one. The contig DN5959_c0_g1_i3 was identified as a defensin named *P. sativum* pI230 mRNA (e-value:0.0, percentage identity: 97.23%). The expression of the DN5959_c0_g1_i3 contig was significantly higher in the tolerant genotypes over the susceptible genotypes under control conditions at 0 (FC=-69.21) and 6 hr (FC=-151.78), and under inoculated conditions at 0 (FC=-7.93) and 12 hr (FC=-43.5). The pI230 mRNA is the precursor for the DRR230 protein, which is a disease resistance response protein identified previously in *P. sativum*. DRR230 defensin was first identified in pea pods in response to infection by *Fsp* (Chiang and Hadwiger, 1991). This gene was found to be overexpressed in pea in the presence of *Micosphaerella pinodes*, a necrotrophic fungus (Fondevilla et al., 2011). This defensin was also found to co-localize with a major QTL (*mpIII-4)* involved in resistance to *M. pinodes* in pea (Prioul-Gervais et al., 2007). DRR230 was isolated by Almeida et al., (2000) and characterized as a small cysteine-rich polypeptide. Almeida et al., (2000) also determined that DRR230 is very effective as a fungal growth inhibitor against *Aspergillus niger, A. vesicolor, Fsph*, and *Neurospora crassa*. While the QTL Fsp-Ps3.3 reported for *Fsp* tolerance (Coyne et al., 2019) is localized on chromosome 5 at 63 million bp position, the location of DRR230 at position 487 million bp makes it an unlikely candidate gene.

The specific function of DRR230 is not yet known, however plant defensins form a characteristic structure known as the cysteine-stabilized α/β motif, a feature that is also shared by several toxins from insects, scorpions, honeybees, and spider venoms (Hadwiger, 2008). Defensins can induce membrane destabilization and inhibit protein synthesis, enzyme activity, and ion channels (Lay and Anderson, 2005; Hadwiger, 2008). The pea DRR230 was overexpressed in canola, and extracts of these plants inhibited the *in vitro* germination of *Leptosphaeria maculans*, a hemibiotrophic fungus (Wang et al., 1999). Canola plants transformed with DRR230 were significantly more resistant to *Leptosphaeria maculans*. The transcriptome analysis presented in this study reinforces recent and preceding studies that suggests that DRR230 may play a key role in resistance or tolerance to *Fsp* induced root rot.

#### 4. Anthocyanin and lignin biosynthetic pathway genes

Pea genotypes with pigmented seed coats and flowers demonstrate highest level of tolerance to *Fsp* (Bodah et al., 2016). It has been hypothesized that genes involved in anthocyanin pigmentation are directly involved in resistance to *Fsp* (Weeden and Porter, 2007; Coyne et al., 2019). However, it was found that the pigmented flowered genotype PI 180693 was susceptible to *Fsp* (Bodah et al., 2016). This result suggests that other genes are potentially involved in the resistance exhibited by pigmented-flowered lines or that several proteins in the anthocyanin pathway could play a role in the production of a colorless metabolite that confers this resistance.

The germplasm utilized in this study consisted of white-flowered lines that present partial tolerance to *Fsp*. RNAseq analysis showed that the white-flowered lines contain a large set of DECs that participate in the anthocyanin biosynthetic pathway (Fig 7). Genes coding for enzymes in the phenylalanine ammonia lyase (PAL) to chalcone isomerase (CHI) biochemical pathway were upregulated in the inoculated treatments in both the tolerant and susceptible genotypes. Furthermore, some of these enzymes were either overexpressed in the susceptible genotypes or were expressed at a similar level between the tolerant and susceptible genotypes (Figure 7).

**Figure 7.**
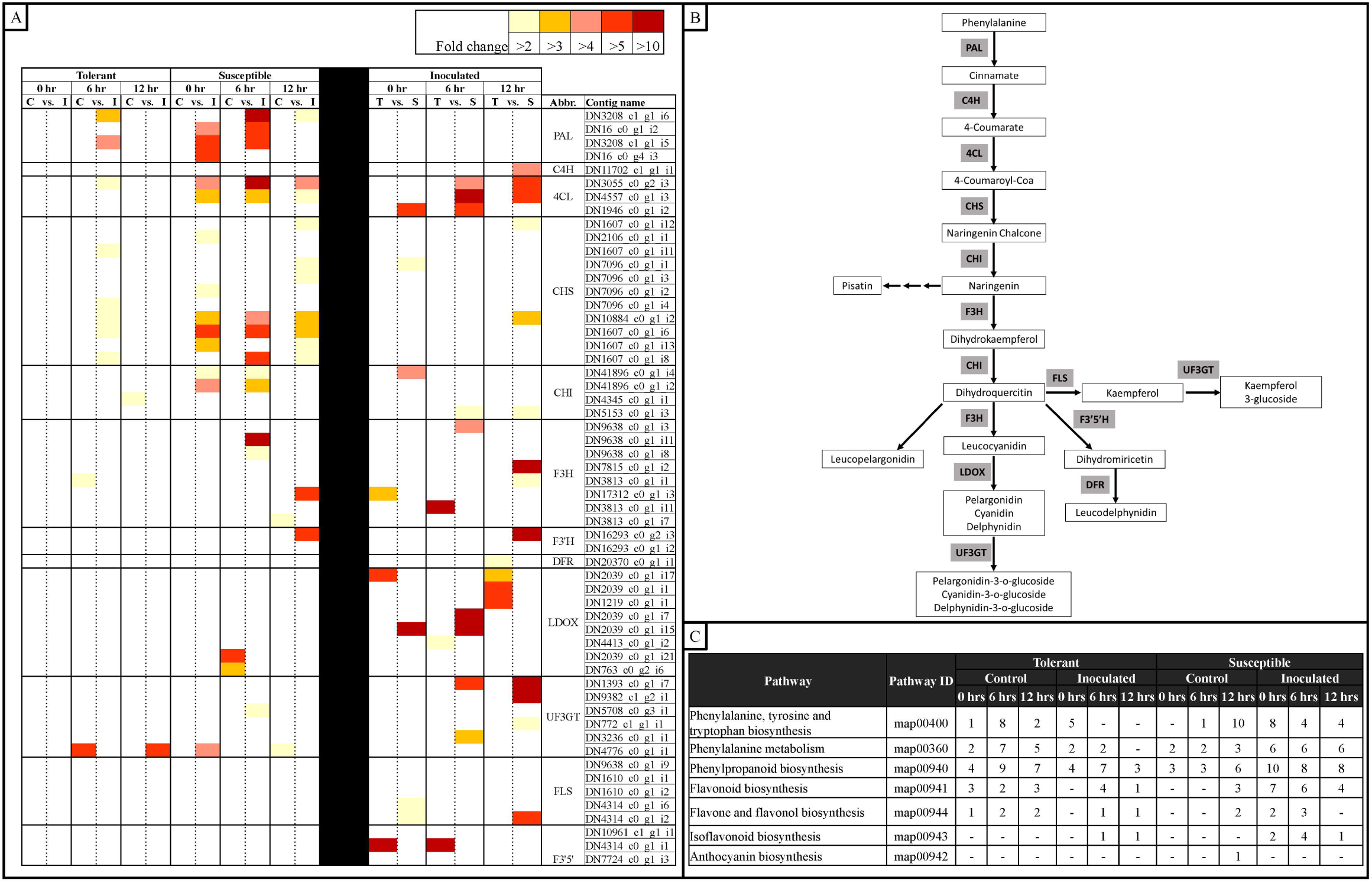
Heatmap representation of changes in the expression of genes associated with the anthocyanin biosynthetic pathway in tolerant and susceptible pea genotypes after *Fsp* challenge. (a) Subset of differentially expressed genes involved in the anthocyanin biosynthetic pathway in tolerant and susceptible pea genotypes after *Fsp* challenge. The color key denotes fold-change. Pairwise comparisons that displayed greater than 2-fold difference (p> 0.005) in expression were identified with a color that ranges from light yellow (fold change>2) to dark red (fold change> 10). (b) Anthocyanin biosynthesis pathway (Adapted from Solfanelli et al., 2006). (c) KEGG pathway analysis of metabolic processes related to the anthocyanin biosynthetic pathway. Abbreviations (Abbr.) PAL= phenylalanine ammonia lyase, CHS= chalcone synthase, CHI= chalcone isomerase, F3H= flavanone-3-hydroxylase, F3’H= flavonoid 3′-hydroxylase, DFR= dihydroflavonol 4-reductase, LDOX= leucoanthocyanidin dioxygenase, UF3GT= UDP glucose-flavonoid 3-o-glucosyl transferase, FLS= Flavonol synthase, F3’5’H= Flavonoid 3′,5′-hydroxylase.

The expression patterns were more variable for genes coding for enzymes from the flavanone-3-hydroxylase (F3H) to LDOX (leucoanthocyanidin dioxygenase), as well as for the flavonoid 3’,5’-hydroxylase (F3’5’), UDP glucose-flavonoid 3-o-glucosyl transferase (UF3GT), and flavonol synthase (FLS) enzymes (Fig 7a, b). Some isoforms of F3H were overexpressed in the susceptible genotypes but also some other isoforms were overexpressed in the tolerant genotypes. F3’H, UF3GT, and FLS were upregulated in the susceptible genotypes, but most or all isoforms of LDOX, and F3’5’ were upregulated in the tolerant genotypes. In the susceptible genotypes, expression of some isoforms of LDOX was suppressed after challenge with *Fsp* (Fig 7a, b).

Coyne et al., (2019) reported a significant QTL *(Fsp-Ps2.1)* that accounts for 44.4 to 53.4% of the phenotypic variance for resistance to *Fsp* and this QTL shows a confidence interval of 1.2 cM. This QTL was found in a population obtained from a cross between a pigmented and a white flower line. *Fsp-Ps2.1* was mapped within the interval of the pigmented flower/anthocyanin pigmentation gene called as gene A in that study. However, the gene was mapped in a white flower cross. One hypothesis is that the resistance gene(s) responsible for the *Fsp-Ps2.1* effect may not necessarily be gene A since *Fsp-Ps2.1* was originally identified in a white-flowered (*a*) cross. The R gene may have been linked in the genome with gene A in the pigmented lines. The white-flowered, resistant parent may have been obtained through a linkage break between *Fsp-Ps2.1* and *A.* Alternatively, a metabolite, possibly a colorless one, in the anthocyanin pathway might be the one that provides this resistance. Fine mapping or gene knockouts are necessary to test this hypothesis.

The transcriptome data, generated in this study, was aligned via BLAST against the *Fsp-Ps2.1* ± 1.2 cM sequence (Table S6). This QTL region was identified in the pea genome (Kreplak et al., 2019) and is approximately 403,600 nt in length. BLAST analysis returned 500 positive blast hits when queried to the transcriptome data. A total of 156 contigs showed differential expression after *Fsp* challenge or when the tolerant and susceptible genotypes expression values are compared (Table S6). A total of 22 of the 156 contigs were annotated as proteins of unknown function or hypothetical proteins. Only the contig TRINITY_DN4823 was identified as a disease related gene, soyasaponin III rhamnosyltransferase. However, this contig was overexpressed in the susceptible genotypes when compared to the tolerant genotypes. No genes associated with pigmentation were identified at the *Fsp-Ps2.1* region during this analysis.

Figure 7c also shows the differential expression of genes at each time point, treatment, and genotype in this study. Early steps in the phenylpropanoid pathway such as phenylalanine biosynthesis, phenylalanine metabolism, phenylpropanoid biosynthesis, flavonoid biosynthesis, flavone and flavonol biosynthesis are active in the control replicates of the tolerant genotypes at the three times points. In the susceptible controls, fewer, or none of the enzymes were overexpressed. This observation suggests that the tolerant genotypes had a higher level of expression of genes in the phenylpropanoid pathway under the basal conditions and, therefore, it was potentially better prepared to defend against *Fsp*.

Legumes contain the isoflavone synthase enzyme, which redirects phenylpropanoid pathway intermediates, such as naringenin, to the synthesis of isoflavonoid phytoalexins (Sreevidya et al., 2006). The isoflavonoid phytoalexins are low molecular weight antimicrobial compounds. Accumulation of these isoflavonoids is often enhanced following infection (Smith and Banks, 1986; Jeandet et al., 2014). Pisatin is an extensively studied phytoalexin from pea. In pea, the presence of *Fsp, Fsph*, and chitosan increases the production of pisatin (Hadwiger and Beckman, 1980). The 6a-hydroxymaackiain-3-O-methyltransferase, enzyme directly upstream from the synthesis of pisatin, was expressed in both the tolerant and susceptible genotypes but overexpressed only in the susceptible genotypes at 0 hrs. Mackintosh et al., (1989) found that the more virulent the *Fsp* isolates are, the more easily they degrade pisatin. The enzyme pisatin demethylase (PDA) demethylates pisatin to produce a less toxic compound. *Fsp* isolates incapable of demethylating pisatin are low in virulence and susceptible to pisatin (Mackintosh et al., 1989; Hadwiger, 2008). Therefore, demethylation of pisatin is an important mechanism by which *Fsp* resists pisatin and a crucial factor in the pathogenicity of *Fsp* in pea. However, based on the results in this and previous studies, pisatin does not seem to play a role in the tolerance to *Fsp* (Mackintosh et al., 1989; Hadwiger, 2008).

The biosynthesis and deposition of lignin in cell walls is developmentally programmed and plays an important role in preventing pathogen invasion (Miedes et al., 2014). The lignin biosynthetic pathway involves the central phenylpropanoid biosynthetic pathway. Genes involved in the lignin biosynthetic pathway, such as PAL, 4CL, trans-cinnamate 4-monooxygenase (C4M) and caffeoyl-o-methyltransferase (COMT), were overexpressed upon *Fsp* inoculation in both genotypes but at a significantly higher level and more consistently in the susceptible genotypes. It is well documented that the lignin biosynthetic pathway produces lignin rapidly in response to cell wall structure perturbations (Caño-Delgado et al., 2003; Tronchet et al., 2010; Sattler and Funnell-Harris, 2013; Miedes et al., 2014). Therefore, it seems the susceptible genotypes are responding to the aggressive *Fsp* invasion with a late and futile effort that involves a higher level of lignin synthesis and deposition.

#### 5. Sugar metabolism

Activation of defense responses upon pathogen infection is usually accompanied by a rapid response in the induction of sink metabolism (Heil and BOSTOCK, 2002; Lanubile et al., 2015). This shift in metabolism is due to the increased demand for carbohydrates as an energy source to sustain the immune responses; however, sugars from the host can also benefit the pathogens in their infection. This study identified DECs involved in sugar transport such as sugar transporter ERD6-like 6, sugar carrier protein C-like, sucrose transport protein SUC3, sugar transport protein 13, probable alkaline/neutral invertase D, bidirectional sugar transporter SWEET2-like, and invertase inhibitor-like protein. These sugars transporters are upregulated in the susceptible genotypes (Table S3). Certain pathogens are known to manipulate plant carbohydrate metabolism for their own needs (Voegele et al., 2001; Lanubile et al., 2015). These pathogens are known to modulate the expression and activity of sugar transporters during their interaction with the plant host. Sugars are used by the pathogen for their own development. Bacterial and fungal pathogens induce the overexpression of different sugar efflux transporters, such as the SWEET genes; this overexpression results in sucrose accumulating in the apoplast for use in pathogen nutritional gain and growth (Chen et al., 2010; Lanubile et al., 2015). Results of this study are in concordance with the literature, suggesting that the active mobilization of sucrose in the *Fsp*-inoculated susceptible genotypes supported successful infection by *Fsp*. Of all the DECs identified in the susceptible genotypes, 78% (25 genes) were overexpressed and only 22% (7 genes) were suppressed after *Fsp* challenge. In the tolerant genotypes, 15% (3 genes) were overexpressed and 85% (17 genes) were suppressed after *Fsp* challenge. These data would support the scenario explained previously; *Fsp* is either manipulating sugar metabolism or taking advantage of the active mobilization of sucrose in the susceptible genotypes.

#### 6. Phytohormones

A large group of DECs were identified that were involved in the synthesis and signaling of salicylic acid (SA), jasmonic acid (JA), and ethylene (ET). The GO enrichment analysis also showed overrepresented GO terms related to the synthesis and signaling of these three hormones. From this set of DECs, a few were overexpressed in the tolerant genotypes, but the vast majority were overexpressed in the susceptible genotypes after *Fsp* challenge.

After pathogen attack, activation of SA- or JA/ET-mediated signaling pathways is known to occur along with induction of expression of pathogenesis-related (*PR*) genes. Some *PR* genes are expressed in response to SA, whereas others in response to JA (Takahashi et al., 2004; Glazebrook, 2005). It is well documented that SA and JA/ET act antagonistically during pathogenesis and other defense-related responses (Clarke et al., 2000; Gupta et al., 2000; Kachroo et al., 2001; Kloek et al., 2001; Shah et al., 2001; Takahashi et al., 2004; Glazebrook, 2005; Leon-Reyes et al., 2010; Shang et al., 2011; Buxdorf et al., 2013). A complex network of interactions among SA, JA, and ET further fine tunes plant defense responses (Feys and Parker, 2000; Kunkel and Brooks, 2002). It is generally assumed that the trophic nature of the pathogen determines which signal transduction pathway (SA or JA/Et) becomes activated in the plant host (Halim et al., 2006). Biotrophic pathogens generally elicit the defense responses via the salicylic acid signaling pathway, while necrotrophs activate a JA-dependent defense response (McDowell and Dangl, 2000; Dangl and Jones, 2001; Thomma et al., 2001; Van Wees et al., 2003; Grant and Lamb, 2006; Halim et al., 2006; Trusov et al., 2009; Rahman et al., 2012).

SA production promotes cell death and that in turn promotes additional SA production. SA-signaling travels through the plant activating systemic acquired resistance (SAR) (Glazebrook, 2005). Necrotrophic pathogens have been shown to hijack plant defense responses to produce SA to further promote disease development. The necrotrophic fungi, *Botrytis cinerea* and *Alternaria solani*, use the SA-signaling pathway to exacerbate the disease in tomato (Rahman et al., 2012). Both pathogens use the SA-signaling pathway through NPR1, a master regulator of SA signaling, and TGA1a TF to promote disease development in tomato. NPR1 and TGA1a suppress the expression of proteinase inhibitors, which in turn suppress the expression of two JA-dependent defense genes (Rahman et al., 2012). The Clover yellow vein virus (ClYVV) also induces cell death via the activation of the salicylic acid (SA) signaling pathway. SA signaling and cell death enhance ClYVV virulence in susceptible pea cultivars (Atsumi et al., 2009).

In this study, differential expression of genes associated with SA synthesis and signaling, cell death and HR in both the tolerant and susceptible genotypes was observed. However, when comparisons were made between the tolerant and susceptible genotypes, these DECs were observed to be overexpressed in the susceptible genotypes. The contig DN5429 in this study was identified as TGA1.a transcription factor (involved in SA signaling) and its isoform DN5429_c0_g1_i1 was overexpressed 6.82-fold in the susceptible genotypes over the tolerant genotypes. These are important observations, at least at the gene expression level, since the current understanding portends that overexpression of SA-related genes should not be observed in interactions between *Fsp* and pea, as cell death in plant hosts does not limit pathogen growth (Glazebrook, 2005). SA-dependent responses and SAR are not predicted to play a role in tolerance against *Fsp*, whereas responses mediated by JA and ET are expected to do so.

The overexpression of genes associated with JA/ET synthesis and signaling also happen in both the tolerant and susceptible genotypes. When comparisons were made between the tolerant and susceptible genotypes, 22 DECs (80% of JA/ET-associated genes) were observed to be overexpressed in the susceptible genotypes. Based on these data, it is difficult to draw conclusions on the effect of SA and/or JA/ET in the response of the tolerant and susceptible genotypes to *Fsp*. However, a comparison was made using the number of DECs associated with the SA and JA/ET biosynthetic and signaling pathway in the tolerant and susceptible genotypes after *Fsp* challenge (Fig 8). The susceptible genotypes showed an upsurge in the overexpression of genes related to SA biosynthesis and signaling (Fig 8). Alternatively, in the tolerant genotypes, the majority of genes related to SA biosynthesis and signaling were suppressed after *Fsp* challenge. These changes related to the SA-pathway genes in the susceptible genotypes might have 1-) deteriorated the action of the JA-signaling pathway, 2-) increased the cell death, and therefore, 3-) facilitated successful infection by *Fsp*

**Figure 8.**
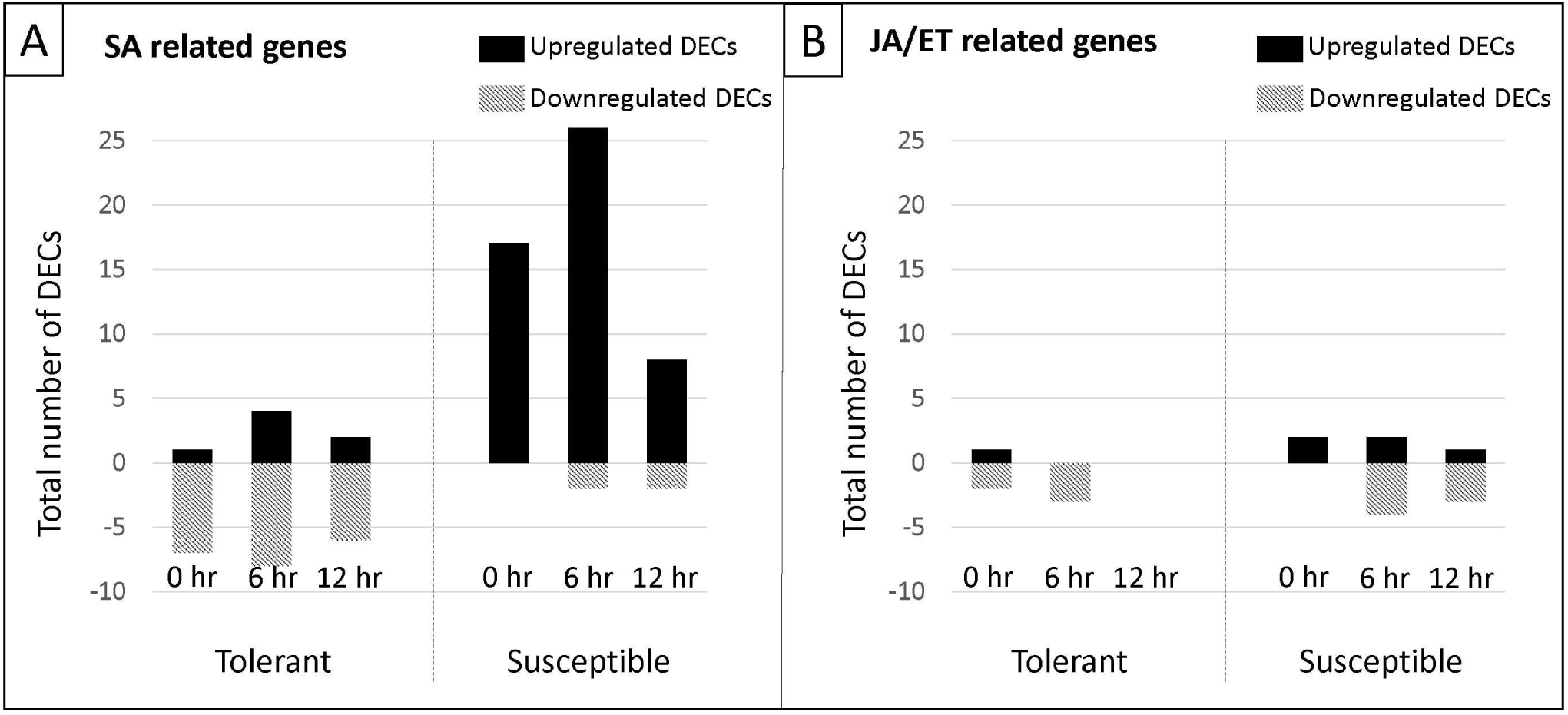
Number of DECs associated with the salicylic (SA) and jasmonate/ethylene (JA/ET) biosynthetic and signaling pathway in a pea tolerant and susceptible genotype after *Fsp* challenge. (a) DECs associated with the salicylic (SA) biosynthetic and signaling pathway. (b) DECs associated with the jasmonate/ethylene (JA/ET) biosynthetic and signaling pathway.

These results suggest that the SA-signaling pathway might be involved in the development of root rot disease caused by *Fsp* in the susceptible genotypes. However, genetic mapping or gene knockouts are needed to evaluate this proposal. Some potential targets for gene knockout experiments would be NPR1 and TGA1.a. Targeting of these genes would provide more insights into the possible manipulation of the SA signaling pathway by *Fsp* in susceptible genotypes.

#### 7. Others: Cell wall and membrane metabolism, and toxin metabolism

Cell wall and membranes play important roles in plant defense as they act as a barrier that prevents pathogen invasion. Both also maintain a reservoir of antimicrobial compounds, which are released during pathogen infection (Vorwerk et al., 2004). Cell wall and membrane modifications are carried out by the plant during pathogen attack. For instance, deposition of callose, which is enriched with Beta-glucan, forms a stronger and thicker barrier at the sites of pathogen attack (Miedes et al., 2014). The formation of this callose appears to be a common mechanism in all plants. However, this response has not been studied in the context of Pea-*Fsp* interaction.

In this study, genes related to cell wall and membrane modification, and callose deposition, were mostly upregulated in the susceptible genotypes (Fig 9).The genes for cell membrane transporters, proteins that work on the detoxification (antiporter activity) of substances, and proteins that break down toxins accumulated in the plant host were also overexpressed in the susceptible genotypes (Fig 9). Interestingly, after *Fsp* challenge, the majority of genes associated with cell wall metabolism, toxin metabolism and transport were suppressed in the tolerant genotypes, while they were overexpressed in the susceptible genotypes (Fig. 9). These responses, at least at the gene expression level, indicate that the response of the susceptible genotypes was delayed as the pathogen had already infested the tissues, and therefore, the host made a futile effort in response to the pathogen attack. Most likely the tolerant genotypes already possessed physical and biochemical barriers and the expression of the genes related to these pathways were actually being suppressed.

**Figure 9.**
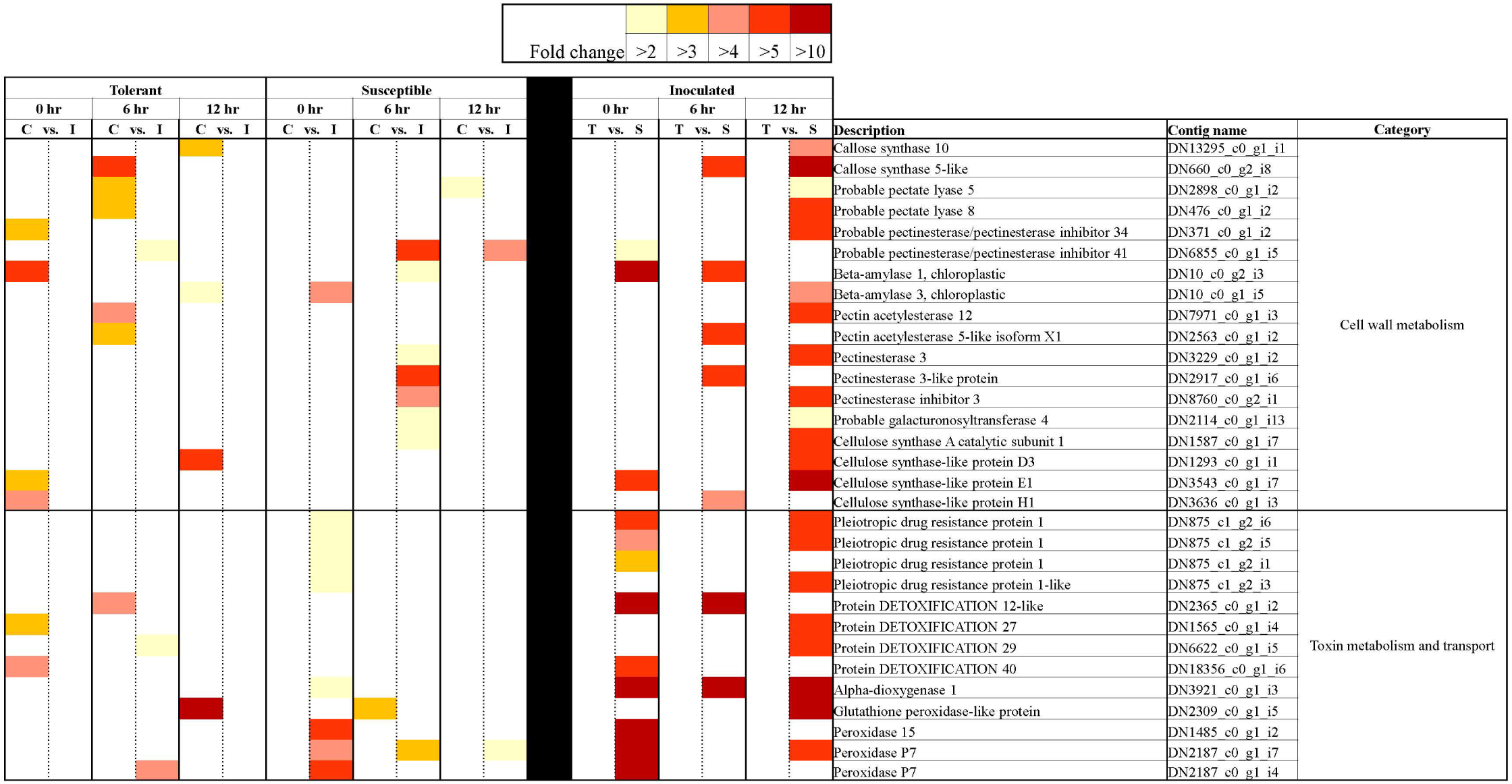
Heatmap representation of differentially expressed genes associated with cell wall metabolism, toxin metabolism and transport in tolerant and susceptible pea genotypes in response to *Fsp* challenge. The color key denotes fold-change. Pairwise comparisons that displayed greater than 2-fold difference (p> 0.005) in expression were identified with a color that ranges from light yellow (fold change>2) to dark red (fold change> 10).

### Conclusions

The time course RNAseq results presented in this study provided a comprehensive insight into the transcriptomic changes that accompany *Fsp* infection in tolerant and susceptible *P. sativum* genotypes. The observed changes in expression of genes are associated with various physiological and biochemical processes that are known to be involved in plant disease response against pathogens. *Fsp* challenge produced a more intense and diverse overexpression of genes, across the entire time-course, in the susceptible genotypes compared to the tolerant genotypes. This type of response is hypothesized to be related to the speed at which the pathogen infestations advances in the susceptible genotypes, and the preexisting level of disease-preparedness in the tolerant genotypes. The transcriptomic effort demonstrated by the susceptible genotypes seems futile and lacked key specific responses that were present in the tolerant genotypes. In contrast, the tolerant genotypes showed a fine-tuned response: fewer changes in the expression of defense-related genes that helps preserve energy, and a faster reset to a basal metabolic state.

This RNAseq analysis helped identify alternate strategies and potential genes that could be evaluated to confer improved tolerance against root rot in *P. sativum*. Specific genes or pathways that might have a key role in tolerance or susceptibility to *Fsp* are: receptor-like cytoplasmic kinase 176, CC-NBS-LRR resistance protein, *WRKY7* TF, WRKY4 TF, *LBD* TF, HMG A TF, anthocyanin biosynthetic pathway, SWEET genes, JA/ET-signaling pathway, cell death, NPR1 and TGA1.a. SA-signaling genes, and most importantly the DRR230 protein. Functional characterization of these genes is expected to provide mechanistic information regarding pea-*Fsp* interaction and provide targets to incorporate resistance via molecular breeding (Bodah et al., 2016) or gene editing (Ghogare et al., 2019) for improving root rot resistance in a crop that is rapidly becoming a meat alternative at the same time as it contributes to sustainability.

## Author Contributions

AD, RS, and BWB designed the study. AD supervised the study. BWB and EB performed the experiments and generated the data. BWB and GN analyzed the data. LP provided the tolerant pea genotypes and *Fsp* isolates. All authors read and approved the final manuscript.

## Acknowledgements

The authors are grateful to Kurt Braunwart, CEO, ProGene Plant Research for critical discussions and his support for the project.

## Data Availability

The raw sequencing data from RNAseq analysis were deposited in the NCBI Sequence Read Archive (SRA, https://www.ncbi.nlm.nih.gov/sra) under the accession number SRP260465.

