## Supplemental Table 1 for "Identification of *Fusarium solani* f. sp. *pisi* (*Fsp*) responsive genes in *Pisum sativum*"

**Table S1. Selected genes, list of primers, annealing temperatures and amplicon sizes for RT-qPCR validation.**

| Contig name | Blast Analysis | | | Primer name | Sequence 5'-->3' | Annealing Temperature (º C) | Size (bp) |
|---|---|---|---|---|---|---|---|
| | Accession | Gene name | E-value | | | | |
| DN2419 | XM_003592027.3 | PREDICTED: Medicago truncatula ethylene-responsive transcription factor ERF110 (LOC11405418), mRNA | 0.0 | BWqPCR_1_fwd | GGAGGAGGATAGCCACCAGT | 64.3 | 122 |
| | | | | BWqPCR_1_rev | CCTGGTCAGCAAATGGAGTT | 64 | |
| DN2754 | XM_004502933.3 | PREDICTED: Cicer arietinum aspartic proteinase Asp1-like (LOC101502257), mRNA | 0.0 | BWqPCR_2_fwd | TCTCGGGTGCTACTGAGAAA | 63.5 | 114 |
| | | | | BWqPCR_2_rev | CCAACCGATGAAGAGTTGGT | 63.9 | |
| DN5727 | MK618561.1 | Lathyrus sativus C2H2-like zinc finger protein mRNA, partial cds | 0.0 | BWqPCR_3_fwd | TGTGCTGTGGAGATGGAAGA | 64.3 | 117 |
| | | | | BWqPCR_3_rev | GGAAAGAGAGGGAAAATCATTG | 62.6 | |
| DN6240 | XM_003592048.3 | PREDICTED: Medicago truncatula uncharacterized LOC11405420 (LOC11405420), mRNA | 0.0 | BWqPCR_4_fwd | GATGGTGAGTGCAAAGCAAA | 63.9 | 143 |
| | | | | BWqPCR_4_rev | AAGCATCAAAAGACTCGGACA | 63.7 | |
| DN2169 | XM_004506541.3 | PREDICTED: Cicer arietinum putative E3 ubiquitin-protein ligase XBAT31 (LOC101496789), mRNAE3 ubiquitin-protein ligase xbat31-like | 6.00E-148 | BWqPCR_5_fwd | TGGAAGGATCACTGGTCACA | 64.4 | 126 |
| | | | | BWqPCR_5_rev | GCAAGTGTAAACGCCATTCA | 63.6 | |
| DN1232 | XM_004504351.3 | PREDICTED: Cicer arietinum stress-associated endoplasmic reticulum protein 2-like (LOC101506542), mRNA | 1.00E-118 | BWqPCR_6_fwd | AAAGGAAAAGACTACCCTGTTGG | 63.2 | 101 |
| | | | | BWqPCR_6_rev | CCACTGGTTGCTGTCCTGAT | 65.2 | |
| DN5529 | XM_024782286.1 | PREDICTED: Medicago truncatula cysteine synthase, chloroplastic/chromoplastic (LOC11443458), mRNA | 2.00E-114 | BWqPCR_7_fwd | TCTCTGGCCCAGTATTCTCA | 62.4 | 111 |
| | | | | BWqPCR_7_rev | CAGTTCGAAAAGCCGAAGAA | 64.2 | |
| DN8631 | XM_013611166.2 | GDP-l-galactose phosphorylase PREDICTED: Medicago truncatula G-type lectin S-receptor-like serine/threonine-protein kinase At1g34300 (LOC25482580), mRNA | 0.0 | BWqPCR_8_fwd | ATTTTGGATTGGCGAAACTG | 63.6 | 120 |
| | | | | BWqPCR_8_rev | CGGCCTTTGAAGTTATTGGA | 63.7 | |
| DN1795 | XM_004514502.3 | PREDICTED: Cicer arietinum GDP-L-galactose phosphorylase 2-like (LOC101497835), mRNA | 0.0 | BWqPCR_9_fwd | TTGCAGTCTTATATTGGGGTTACA | 63.1 | 100 |
| | | | | BWqPCR_9_rev | GATCTACACGGCCTCATTGC | 64.5 | |
| DN8314 | AF139187.1 | Pisum sativum root border cell-specific protein (BRD13) mRNA, complete cds | 0.0 | BWqPCR_Ctrl_fwd | TGCAGTTGCTGAAGTGTTCC | 64.1 | 128 |
| | | | | BWqPCR_Ctrl_rev | GTAACGAAGGGTTGCGTGAT | 63.8 | |
