## Supplemental Table 2 for "Identification of *Fusarium solani* f. sp. *pisi* (*Fsp*) responsive genes in *Pisum sativum*"

**Table S2. Summary of Illumina reads and mapping results from RNA-seq analysis for each library generated in this study.**

| Library ID | Genotype (bulk) | Condition | Biological replicate | Number of raw reads | mean _Q | Number of reads after trimmed | Percentage trimmed (%) | Contig genes mapped | Contig genes mapped percentage (%) | Average length of contigs |
|---|---|---|---|---|---|---|---|---|---|---|
| P.sativum_S1 | Tolerant | 0hr Control | 1 | 21,087,280 | 34.69 | 14,668,172 | 30.44 | 125,240 | 67.44 | 1,526.30 |
| P.sativum_S2 | | | 2 | 29,751,755 | 34.49 | 20,500,575 | 31.09 | | | |
| P.sativum_S5 | | 0hr Inoculated | 1 | 17,591,660 | 34.22 | 12,025,245 | 31.64 | 103,377 | 55.66 | 1,651.56 |
| P.sativum_S6 | | | 2 | 38,325,318 | 34.31 | 26,156,099 | 31.75 | | | |
| P.sativum_S9 | | 6hr Control | 1 | 24,060,774 | 34.23 | 16,316,996 | 32.18 | 127,056 | 68.41 | 1,561.63 |
| P.sativum_S10 | | | 2 | 36,645,962 | 34.68 | 25,483,911 | 30.46 | | | |
| P.sativum_S13 | | 6hr Inoculated | 1 | 33,314,160 | 34.35 | 22,806,162 | 31.54 | 110,964 | 59.74 | 1,665.00 |
| P.sativum_S14 | | | 2 | 45,975,152 | 34.50 | 31,791,950 | 30.85 | | | |
| P.sativum_S17 | | 12hr Control | 1 | 30,568,144 | 34.57 | 21,079,674 | 31.04 | 126,569 | 68.15 | 1,564.87 |
| P.sativum_S18 | | | 2 | 37,766,216 | 34.54 | 26,055,362 | 31.01 | | | |
| P.sativum_S21 | | 12hr Inoculated | 1 | 38,657,700 | 34.50 | 26,658,083 | 31.04 | 102,382 | 55.13 | 1,738.31 |
| P.sativum_S22 | | | 2 | 21,378,038 | 34.26 | 18,145,808 | 15.12 | | | |
| P.sativum_S3 | Susceptible | 0hr Control | 1 | 22,860,375 | 34.12 | 16,910,696 | 26.03 | 123,269 | 66.37 | 1,532.24 |
| P.sativum_S4 | | | 2 | 21,695,910 | 34.48 | 14,958,281 | 31.05 | | | |
| P.sativum_S7 | | 0hr Inoculated | 1 | 35,432,148 | 34.45 | 24,358,895 | 31.25 | 137,378 | 73.97 | 1,449.52 |
| P.sativum_S8 | | | 2 | 40,203,272 | 34.52 | 27,819,393 | 30.80 | | | |
| P.sativum_S11 | | 6hr Control | 1 | 35,785,070 | 34.88 | 25,115,168 | 29.82 | 134,800 | 72.58 | 1,506.69 |
| P.sativum_S12 | | | 2 | 27,813,826 | 34.43 | 19,097,253 | 31.34 | | | |
| P.sativum_S15 | | 6hr Inoculated | 1 | 42,437,824 | 34.28 | 28,918,745 | 31.86 | 140,566 | 75.69 | 1,446.26 |
| P.sativum_S16 | | | 2 | 39,370,422 | 34.00 | 26,476,697 | 32.75 | | | |
| P.sativum_S19 | | 12hr Control | 1 | 63,685,406 | 34.62 | 44,134,320 | 30.70 | 141,530 | 76.21 | 1,467.68 |
| P.sativum_S20 | | | 2 | 56,070,996 | 34.71 | 38,889,470 | 30.64 | | | |
| P.sativum_S23 | | 12hr Inoculated | 1 | 34,620,388 | 34.86 | 24,188,558 | 30.13 | 132,009 | 71.08 | 1,531.62 |
| P.sativum_S24 | | | 2 | 50,891,022 | 34.75 | 35,387,679 | 30.46 | | | |
