## Supplemental Table 4 for "Identification of *Fusarium solani* f. sp. *pisi* (*Fsp*) responsive genes in *Pisum sativum*"

**Table S4. Pairwise comparisons of expression values (read counts) for each contig to obtain differentially expressed contigs (DECs).**

| Comparison | Group 1 | | | Group 2 | | |
|---|---|---|---|---|---|---|
| | Genotype | Time point (hr) | *Fsp* | Genotype | Time point (hr) | *Fsp* |
| 1 | Tolerant | 0 | Control | Tolerant | 0 | Inoculated |
| 2 | Tolerant | 6 | Control | Tolerant | 6 | Inoculated |
| 3 | Tolerant | 12 | Control | Tolerant | 12 | Inoculated |
| 4 | Susceptible | 0 | Control | Susceptible | 0 | Inoculated |
| 5 | Susceptible | 6 | Control | Susceptible | 6 | Inoculated |
| 6 | Susceptible | 12 | Control | Susceptible | 12 | Inoculated |
| 7 | Tolerant | 0 | Control | Susceptible | 0 | Control |
| 8 | Tolerant | 6 | Control | Susceptible | 6 | Control |
| 9 | Tolerant | 12 | Control | Susceptible | 12 | Control |
| 10 | Tolerant | 0 | Inoculated | Susceptible | 0 | Inoculated |
| 11 | Tolerant | 6 | Inoculated | Susceptible | 6 | Inoculated |
| 12 | Tolerant | 12 | Inoculated | Susceptible | 12 | Inoculated |
